# Optogenetic inhibition of light-captured alcohol-taking striatal engrams facilitates extinction and suppresses reinstatement

**DOI:** 10.1101/2024.06.02.597021

**Authors:** Valerie Vierkant, Xueyi Xie, Zhenbo Huang, Lian He, Eric Bancroft, Xuehua Wang, Rahul Srinivisan, Yubin Zhou, Jun Wang

**Author notes:** These authors contributed equally (Valerie Vierkant, Xueyi Xie). To whom correspondence should be addressed: Jun Wang, M.D., Ph.D., 8447 Riverside Pkwy, 2106 Medical Research and Education Building Bryan, TX 77807.

## Abstract

**Background:** Alcohol use disorder (AUD) is a complex condition, and it remains unclear which specific neuronal substrates mediate alcohol-seeking and -taking behaviors. Engram cells and their related ensembles, which encode learning and memory, may play a role in this process. We aimed to assess the precise neural substrates underlying alcohol-seeking and -taking behaviors and determine how they may affect one another.

**Methods:** Using FLiCRE (Fast Light and Calcium-Regulated Expression; a newly developed technique which permits the trapping of acutely activated neuronal ensembles) and operant-self administration (OSA), we tagged striatal neurons activated during alcohol-taking behaviors. We used FLiCRE to express an inhibitory halorhodopsin in alcohol-taking neurons, permitting loss-of-function manipulations.

**Results:** We found that the inhibition of OSA-tagged alcohol-taking neurons decreased both alcohol-seeking and -taking behaviors in future OSA trials. In addition, optogenetic inhibition of these OSA-tagged alcohol-taking neurons during extinction training facilitated the extinction of alcohol-seeking behaviors. Furthermore, inhibition of these OSA-tagged alcohol-taking neurons suppressed the reinstatement of alcohol-seeking behaviors, but, interestingly, it did not significantly suppress alcohol-taking behaviors during reinstatement.

**Conclusions:** Our findings suggest that alcohol-taking neurons are crucial for future alcohol-seeking behaviors during extinction and reinstatement. These results may help in the development of new therapeutic approaches to enhance extinction and suppress relapse in individuals with AUD.

## INTRODUCTION

Alcohol use disorder (AUD) is a chronic disease characterized by compulsive alcohol seeking and consumption (Carvalho et al., 2019, Kranzler and Soyka, 2018). Over 22,000 people aged 26 and up were diagnosed with AUD in 2020, and an estimated 95,000 people die from alcohol-related causes annually (National Center and Drug Abuse, 2023). Despite the severity of this disorder, currently developed treatment methods have only limited effects (Ray et al., 2021). Additionally, even following treatment, there is a large rate of individuals who relapse and return to drinking (Dawson et al., 2007). This return to the seeking and consumption of alcohol remains a major barrier to the treatment of AUD and is likely in part due to long-lasting neurochemical changes that occur with exposure to alcohol (Stewart, 2008, Sliedrecht et al., 2019). Subsequently, research into the neural mechanisms underlying relapse to alcohol drinking is important for the future development of effective treatments.

AUD has been previously characterized as a maladaptive form of learning and memory (Hyman et al., 2006, Milton and Everitt, 2012). Memories are thought to be encoded by populations of cells in various brain regions that are physically and chemically modified by the learning experience, collectively forming what is termed engram cells (Josselyn and Tonegawa, 2020). Engram cell ensembles are thought to contain engram cells that contribute to the associated memories (Sweis et al., 2021). Several ensembles have been identified that are related to AUD-associated memories (George and Hope, 2017, Visser et al., 2020). For example, ensembles located within the infralimbic cortex are thought to inhibit cue-induced alcohol-seeking (Pfarr et al., 2015), whereas the recruitment of an ensemble located within the central nucleus of the amygdala during withdrawal is thought to promote excessive alcohol consumption behaviors (de Guglielmo et al., 2016). Despite these findings, current treatments for AUD lack specificity on the ensembles driving alcohol-associated behaviors, which may contribute to their limitations.

Thus, a better understanding of the ensembles underlying alcohol-related behaviors can promote future development of more specific treatments.

AUD is made up of two critical behaviors: alcohol seeking and alcohol taking (Lüscher et al., 2020). Several studies have used optogenetic or chemogenetic techniques to study the behaviors underlying AUD (Hellard et al., 2019, Cheng and Wang, 2019, Cheng et al., 2021), but few have examined the interplay of neural substrates related to these distinct behaviors. However, there is evidence to believe that alcohol-seeking and -taking behaviors may involve distinct cellular substrates (Lüscher et al., 2020); for example, Renteria et al. (2020) found no relationship between the degree of habitual EtOH seeking and escalated consumption. In addition, another study found that antagonism of α4β2 and α3β4 nicotinic acetylcholine receptors had different effects on taking and seeking behaviors (Cippitelli et al., 2018). However, it is still unknown for certain if alcohol-seeking and -taking behaviors are mediated by the same or different neural ensembles, or how the ensembles for one behavior might affect the other.

This study aimed to determine if distinct neural substrates mediate alcohol-seeking and – taking behaviors and determine how they contribute to the extinction and reinstatement of alcohol-seeking and -taking behaviors. Specifically, we focused on the dorsomedial striatum (DMS), a brain region critically implicated in driving goal-directed behaviors. The DMS has been shown to play a crucial role in AUD and in mediating its related behaviors (Wang et al., 2015, Roltsch Hellard et al., 2019), and is thus a relevant target for dissecting the two target behaviors.

The lack of literature focusing on the difference between seeking and taking behaviors in AUD most likely arises due to the short timeframe that occurs between the two behaviors. This prevents the use of standard activity-dependent labeling technologies, such as FosTRAP and ArcTRAP, which can take hours to use. Thus, this study utilized a novel technology called FLiCRE (Fast Light- and Calcium-Regulated Expression), which enables the characterization of acutely-activated neuronal ensembles with a minute-timescale sensitivity (Kim et al., 2020). This method permits the expression of a transcription factor with activity gated by both blue light and calcium to trap activated neurons. These transcription factors can later be manipulated within the trapped neurons for gain or loss of function studies.

This study utilized alcohol operant self-administration (OSA) and FLiCRE in D1-Cre rats to capture alcohol-taking (magazine entry) associated neurons in the DMS in order to test their role in future alcohol-taking and -seeking behaviors. Alcohol OSA was used to measure both alcohol-seeking and -taking behaviors using lever presses and magazine entries, respectively (Blegen et al., 2018), and FLiCRE was used to express an inhibitory halorhodpsin (eNphR; Gradinaru et al., 2008) in the trapped alcohol-taking neurons. Thus, yellow light could be used to inhibit alcohol-taking neurons in future manipulations. We first tested the effects of their inhibition during regular OSA, and then assessed the role of OSA-tagged magazine entry neurons in extinction and reinstatement.

In the present study, we demonstrated that optogenetic inhibition of magazine entry striatal ensembles during OSA reduced both seeking and taking behaviors. Additionally, we found that the suppression of OSA-tagged alcohol-taking neurons during extinction training facilitated the extinction of alcohol-seeking behaviors. Finally, we showed that subsequent inhibition of these ensembles suppressed the reinstatement of alcohol seeking. Our data suggests that alcohol-taking neurons are essential for future alcohol-seeking behaviors during extinction and reinstatement.

## MATERIALS AND METHODS

### Experimental model and subject details Animals

D1-Cre rats were obtained from the NIDA. Wild-type (WT) C57BL/6J mice were purchased from the Mutant Mouse Regional Resource Centers. Genotyping was performed using PCR of tail DNA. Both male and female transgenic littermates (3–4 months old) were used in this study. Animals were housed with 2–4 gender-matched conspecifics in a 12-h light-dark cycle (light from 7 a.m. to 7 p.m.) with *ad libitum* food and water. The room temperature for all animal housing and experiments was 23°C. Investigators were blind to all experimental groups. All animal care and experimental procedures were approved by the Texas A&M University Institutional Animal Care and Use Committee and were conducted in agreement with the National Research Council Guide for the Care and Use of Laboratory Animals.

### Primary Neuronal Culture and Packing AAV

All adeno-associated viruses (AAVs) were packaged in house and the packing protocol was adapted from Feng Zhang’s lab at MIT and Alice Ting’s lab at Stanford (Kim et al., 2020).

#### HEK293 cells

HEK293T cells were grown as a monolayer in complete Dulbecco’s Modified Eagle Medium (DMEM, Corning) supplemented with 1% penicillin-streptomycin (Corning, containing 5000 units/mL penicillin and 5000 mg/mL streptomycin) and 10% Fetal Bovine Serum (FBS, VWR) at 37°C with 5% CO2. The cell line has not undergone authentication.

#### AAV1/2 production

AAV1/2 viruses were prepared and concentrated as previously reported (Wang et al., 2017, Kim et al., 2020), and the steps are briefly summarized here. HEK293T cells were grown to 90% confluency (described above) in 3-4 T150 flasks per construct, and then each flask was transfected with 5.2 ug vector of interest, 4.35 ug AAV1 plasmid, 4.35 µg AAV2 plasmid, 10.4 µg DF6 AAV helper plasmid, and 130 ml PEI. Cells underwent incubation for 48 hr. Following incubation, supernatant media was collected to infect cultured neurons. The remaining HEK293T cells were pelleted at 800 g for 10 min and subsequently resuspended in 20 mL of 150 mM TBS (150 mM NaCl, 20 mM Tris, pH = 8.0) for purification and concentration. The resuspended cells then had 10% sodium deoxycholate (in water, Sigma) added at a final concentration of 0.5%. Additionally, 50 units/mL of benzonase nuclease (Sigma) was added. The cell suspension was incubated for 1 hour at 37°C. Following incubation, it was cleared by 15 min of centrifugation at 3,000 g. A peristaltic pump (MPII, Harvard Apparatus) was used to load the supernatant to a HiTrap heparin column (GE Healthcare), pre-equilibrated with 10 mL of TBS. Then, 20 mL of 100 mM TBS was used to wash the column using the pump, followed by using a syringe to wash with 1 mL of 200 mM TBS and 1 mL of 300 mM TBS. Subsequently, 1.5 mL 400 mM TBS, 3.0 mL 450 mM TBS, and then 1.5 mL of 500 mM TBS were used to elute the virus, and then a 15 mL centrifugal unit (100K molecular weight cut-off, Amicon) was used for concentrating the virus, spinning at 2,000 g for ∼1 min to a final volume of 500 µL. A 1.5 mL centrifugal unit (Amicon) was used for further concentrating the virus to ∼100 ml. Then, 2 ml of virus was incubated with 1 ml DNase I (NEB, 2 units/ml), 4 µL DNase I buffer, and 33 ml H2O for 30 min at 37°C to titer the concentrated viruses. The DNase I was inactivated at 75°C for 15 min, and then 5 ml of this reaction was added to 14 ml H2O and 1 µL of Proteinase K (Thermo Fisher, 20 mg/mL) at 50°C for 30 min. Subsequently, proteinase K was inactivated for 10 min at 98°C. For the qPCR reaction, 2 ml of this reaction, along with 5 µL of SYBR Green master mix (2x), 0.06 ml each of the forward and reverse primers (0.3 mM), and 2.88 ml of H2O, were used. The primers were designed against the synapsin promoter and WPRE (see (Kim et al., 2020) for details). Purified/linearized AAV DNA plasmids containing synapsin and WPRE were used to generate standardized curves, with three different standard curves generated for 0.05 ng, 0.1 ng, or 0.2 ng of linearized DNA. Each viral sample titer was calculated in reference to the standard curve as previously described (Wang et al., 2017).

### Surgery

#### Stereotaxic virus infusion and fiber implantation

Virus infusion was conducted as described previously (Lu et al., 2021, Cheng et al., 2021, Ma et al., 2021). Animals were anesthetized with 3–4% isoflurane at 1.0 L/min and immobilized in a stereotaxic surgery frame with ear bars (David Kopf Instruments); lubricant ophthalmic ointment was applied to prevent eye drying. After a midline scalp incision, small bilateral craniotomies were made using a microdrill (0.5 mm burr), based on the mouse brain atlas. For AAV validation, AAV-CaM-GFP, AAV-LOV, and AAV-TRE-mCherry were infused into the DMS of WT mice (AP: +0.38 mm, ML: ±1.8 mm, DV: −2.9 mm from the Bregma) using coordinates adapted from previous research (Lu et al., 2019) at a rate of 0.12 µL/min (2.1 µL/site, 0.7 µL of each virus, bilaterally). For behavioral manipulation, AAV-CaM-GFP, AAV-LOV, and AAV-TRE-eNphR-EYFP were infused into the DMS of D1-Cre rats (AP: +0.36 mm, ML: ±2.4 mm, DV: −4.8 mm from the Bregma) using coordinates adapted from previous research (Ma et al., 2018) at a rate of 0.12 µL/min (1.3 µL/site, 0.3 µL AAV-CaM-GFP, 0.5 µL AAV-loV, and 0.5 µL AAV-TRE-eNphR-EYFP). The injectors were maintained at the injection sites for 10 min after the end of the infusion to allow virus diffusion. Fibers (200-μm diameter optical fiber secured to a 1.25-mm stainless steel ferrule for mice; 300-μm diameter optical fiber secured to a 2.5-mm stainless steel ferrule for rats) were bilaterally implanted through the injection site, with DV 0.1 mm above the virus infusion. For fiber implants, four metal screws were fixed into the skull to support the implants, which were further secured with dental cement (Henry Schein). After the surgery, the animals were returned to their home cages for at least 6 weeks for recovery and virus expression.

#### AAV in vivo validation

Three weeks after the surgery, we anesthetized the WT mice using isofluorane. We then separated the WT mice into three groups: one group received cocaine injection (i.p., 15 mg/kg) without blue light stimulation, one group received blue light stimulation without neuronal activation from cocaine, and one group received both cocaine injection (i.p., 15 mg/kg) and blue light stimulation. For the stimulation protocol, we used 20 Hz, 10-ms pulse width ∼5mW power measured at the patch cable. The stimulation was delivered as 1 min on, 2 min off, for a total duration of 15 min (Kim et al., 2020). Three weeks after the stimulation, we perfused animals to conduct confocal scanning.

### Electrophysiological recordings of brain slices

Brain slices were prepared as previously described (Lu et al., 2019, Ma et al., 2021, Ma et al., 2018). Rats were anesthetized with isoflurane and underwent cardiac perfusion with ice-cold cutting solution. The brains were removed rapidly and coronal slices (250 µm in thickness) of the striatum were cut in an ice-cold cutting solution containing the following (in mM): 40 NaCl, 143.5 sucrose, 4 KCl, 1.25 NaH_2_PO_4_, 26 NaHCO_3_, 0.5 CaCl_2_, 7 MgCl_2_, 10 glucose, 1 sodium ascorbate and 3 sodium pyruvate (pH 7.35, 305–310 mOsm), saturated with 95% O_2_ and 5% CO_2_. Slices were then incubated in a 1:1 mixture of cutting solution and external solution at 32°C for 45 min. The external solution was composed of the following (in mM): 125 NaCl, 4.5 KCl, 2 CaCl_2_, 1 MgCl_2_, 1.25 NaH_2_PO_4_, 25 NaHCO_3_, 15 sucrose and 15 glucose, pH 7.35, 305–310 mOsm), saturated with 95% O_2_ and 5% CO_2_. Slices were maintained in an external solution at room temperature until use. Recordings were performed as previously described (Lu et al., 2019, Ma et al., 2021, Cheng et al., 2018, Ma et al., 2018, Huang et al., 2017). The brain slice was positioned in a recording chamber attached to the fixed stage of an upright microscope (Olympus) and perfused with an oxygenated external solution at 32°C, with a flow rate of 2 mL/min. Neurons were visualized using a 40× water-immersion lens and an infrared-sensitive CCD camera. A Multiclamp 700B amplifier with Clampex 10.6 software and Digidata1550 data acquisition system (Molecular Devices, Sunnyvale, CA) were used for recording. A recording pipette with a resistance of 3–6 MΩ was pulled from a borosilicate glass capillary (World Precision Instruments, Sarasota, FL) using a micropipette puller (Model P-97, Sutter Instrument Co.). Recording electrodes contained internal solution (in mM: 125 CsMeSO4, 6 NaCl, 10 HEPES, 1 EGTA, 10 QX-314.Cl, 2 MgATP, 6 Na_3_GTP, 2 Na_2_CrPO_4_; pH 7.25, 280 mOsm). To measure optogenetically induced inhibitory currents in TRE-eNphR-eYFP expressed neurons, whole-cell voltage-clamp recordings were used. eYFP-positive neurons were clamped at 0 mV and 590 nm light of different durations were delivered to brain slices through an objective lens.

### Behavioral testing

#### Operant Self Administration

D1-Cre rats were trained to self-administer a 20% alcohol (v/v) in water in an OSA chamber as previously described (Lu et al., 2019, Ma et al., 2021, Cheng et al., 2018, Ma et al., 2018, Huang et al., 2017). Each chamber contains two levers: pressing the active lever results in the delivery of 0.1 ml of the alcohol solution; pressing of the inactive lever is recorded, but no programmed events occur. The alcohol solution was delivered by a stainless-steel dipper that rested within an alcohol reservoir. When activated, the dipper was raised into the magazine port for 20 sec, then returned to the reservoir. The weight of the alcohol reservoir was measured before and after the operant session to determine how much alcohol a rat consumed during the operant session. During alcohol delivery, the magazine port was illuminated, which served as a cue to signal reward availability. We started training with a Fixed Ratio 1 (FR1) schedule, where one active press resulted in a delivery of 0.1 ml of the alcohol solution. The schedule of delivery was gradually increased up to a Fixed Ratio 7 (FR7) schedule, where seven active presses resulted in a delivery of 0.1 ml of the alcohol solution. Animals were trained on this schedule until they reached steady responding. Following this, we captured active neurons using blue light across three consecutive sessions. During these sessions, blue light was delivered contingently with each alcohol delivery for the first ten seconds. After capturing, we maintained animals on the FR7 schedule for three weeks prior to any manipulation. For optogenetic inhibition, we delivered yellow light (5 mW; 2 sec on, 2 sec off) throughout the entire operant sessions.

#### Extinction Training

Animals underwent 9 sessions of extinction training, in which active presses no longer resulted in alcohol delivery or cue light presentation. For the experimental group, yellow light stimulation was delivered (5 mW; 2 sec on, 2 sec off) for the first four sessions of extinction training. For the control group, no yellow light stimulation was performed. We removed day one of extinction due to additional stress from habituation to the light and poor extinction responding from it being the first day of extinction.

#### Reinstatement

The reinstatement test was conducted as described elsewhere (Wang et al., 2010). It was carried out at the FR7 schedule, in which seven presses on the active side triggered the illumination of the cue light within the magazine port. However, there was an exception wherein the first cue light was delivered non-contingently at the beginning of the session. Since following the 9-day extinction training multiple animals stopped active responses, 20 µL of alcohol was pipetted into the magazine port before the session’s start to instigate responses from the animals. However, once the session had started, alcohol was not available. For the experimental group, yellow light stimulation was delivered (5 mW; 2 sec on, 2 sec off) for the first four sessions of extinction training. For the control group, no yellow light stimulation was performed.

### Histology

Histology studies for the FLiCRE AAV validation were performed as described previously (Wei et al., 2018, Lu et al., 2021). The mice were first perfused with PBS and then fixed with 4% paraformaldehyde, following which the brains were removed and post-fixed in 4% paraformaldehyde for 24 h, dehydrated with 30% sucrose for 3 d at 4°C, and coronally cut into 50-μm slices at −22°C. VECTASHIELD mounting media (Vector Laboratories) was used to mount the slices on slides after they were washed in PBS. Confocal microscopy (Fluoroview-1200, Olympus) was used to obtain images which were then analyzed by Imaris 8.3.1 (Bitplace, Zurich, Switzerland). Identical parameters were used for imaging and analysis of all specimens.

### Quantification and Statistical analysis

Electrophysiological data analyses were conducted with both the Clampfit software (Molecular Devices, Sunnyvale, CA) and the Mini Analysis Program (Synaptosoft, Decatur, GA). SigmaPlot 12.5 (Systat Software Inc.) and Prism (Graphpad) were used for all statistical analyses. Experiments with more than two groups were subjected to one-way ANOVA, two-way ANOVA, one-way repeated measures ANOVA (RM ANOVA), or two-way RM ANOVA with Turkey *post-hoc* tests for multiple comparisons. Two-tailed paired or unpaired Student’s *t* tests were used to analyze experiments with two groups. All data are presented as the mean ± s.e.m.

## RESULTS

### FLiCRE expression is both light- and activity-dependent

We first validated the use of FLiCRE in WT mice. We infused the AAVs encoding FLiCRE (CaM-GFP, LOV, and TRE-mCherry) into the DMS (Fig. 1A). In this system, calcium influx induced by neuronal activity will lead to calmodulin (CaM)-GFP carrying TEV protease binding to its partner in the light-oxygen-voltage domain (LOV) complex, permitting the protease to be close to the protease cleavage site covered by the LOV complex. At the same time, the delivery of blue light will activate LOV to expose the protease cleavage site, allowing the protease in the CaM-GFP complex to cleave and release tetracycline-controlled transactivator protein (tTA). This tTA will translocate into the nucleus to activate the transcription of TRE-mCherry. TRE-mCherry was selected as the transcription factor so that tagged neurons would express mCherry and be identifiable to verify that the method was working as intended. Optical fibers were also implanted into the DMS to permit blue light delivery (Fig. 1B).

**Figure 1.**
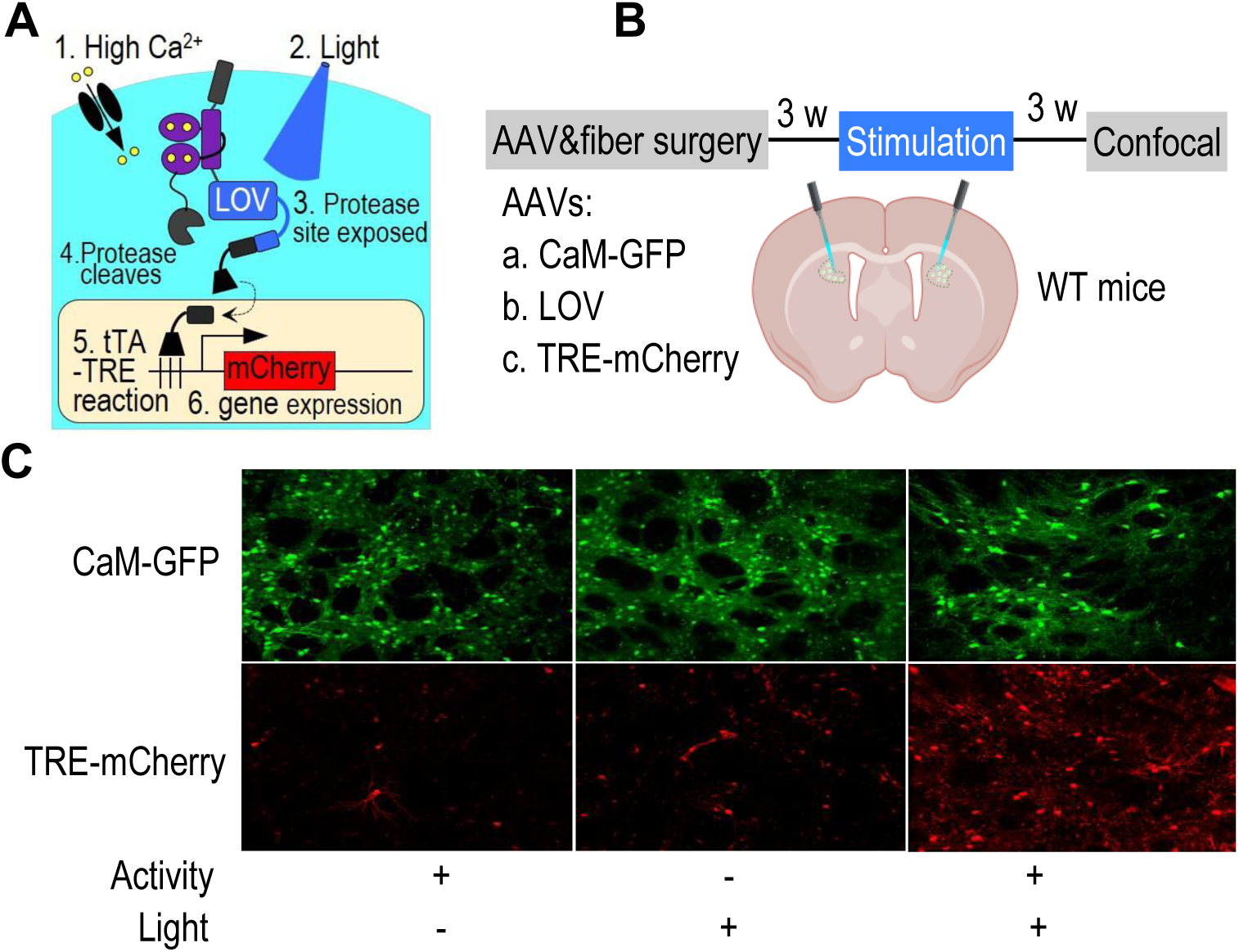
FLiCRE expression is both light- and activity-dependent. ***A***, Schematic of FLiCRE-driven mCherry expression. Intracellular calcium rise (1) and light stimulation of LOV (2) triggers protease site exposure (3) and enables tTA cleavage (4). tTA translocates to the nucleus (5), activating mCherry expression (6). LOV, light-oxygen-voltage domain (open by blue light). ***B***, Timeline of validation experiment. Three weeks after AAV infusion and fiber implantation into the DMS of WT mice, animals were intraperitonially injected with cocaine (15 mg/kg) and exposed to blue light stimulation. After an additional three weeks, mice were perfused for confocal imaging. ***C***, Confocal images showing activity- and light-dependent mCherry expression. Note that CaM-GFP is constant across all conditions, but the highest expression of mCherry only occurs when neuronal activity (induced by cocaine injection) and light stimulation are paired.

Three weeks after the virus infusion, neuronal activity following cocaine exposure *in vivo* and blue light were used to trap active neurons. Three different groups were used (Fig. 1C); one group had neurons activated via cocaine exposure but did not receive blue light, one group received blue light without neuronal activation, and the final group had active neurons and received blue light. We found that only the group with both neuronal activity and blue light administration showed high levels of mCherry expression (Fig. 1C), validating that FLiCRE is both light- and activity-dependent.

### Inhibition of magazine-entry-tagged neurons decreased alcohol-taking behaviors

Following the validation of using FLiCRE to tag acutely activated neuronal ensembles, we infused AAVs containing FLiCRE (CaM-GFP, LOV, and TRE-eNphR-eYFP) that would drive the expression of the inhibitory halorhodopsin eNphR and implanted optical fibers into the DMS of D1-Cre rats. The rats were trained on alcohol OSA until they reached steady responding on an FR7 schedule. To capture the neurons active during magazine entry, blue light was delivered for 10 seconds upon the start of magazine entry (Fig. 2A). Although the delivery period is 20 seconds, we used blue light stimulation during only the first 10 seconds as it takes around 10 seconds following the removal of the blue light for the protease response to turn off (Kim et al., 2020). This blue light stimulation was repeated throughout multiple trials of each session; sessions were made up of around 70 trials, suggesting that there were 700 seconds (around 12 minutes) of blue light stimulation for tagging neurons that were active during magazine entries during each session. This method for tagging active neurons is consistent with previous research (Lee et al., 2017). To ensure tagging of a sufficient number of neurons, we delivered blue light for 3 consecutive sessions.

**Figure 2.**
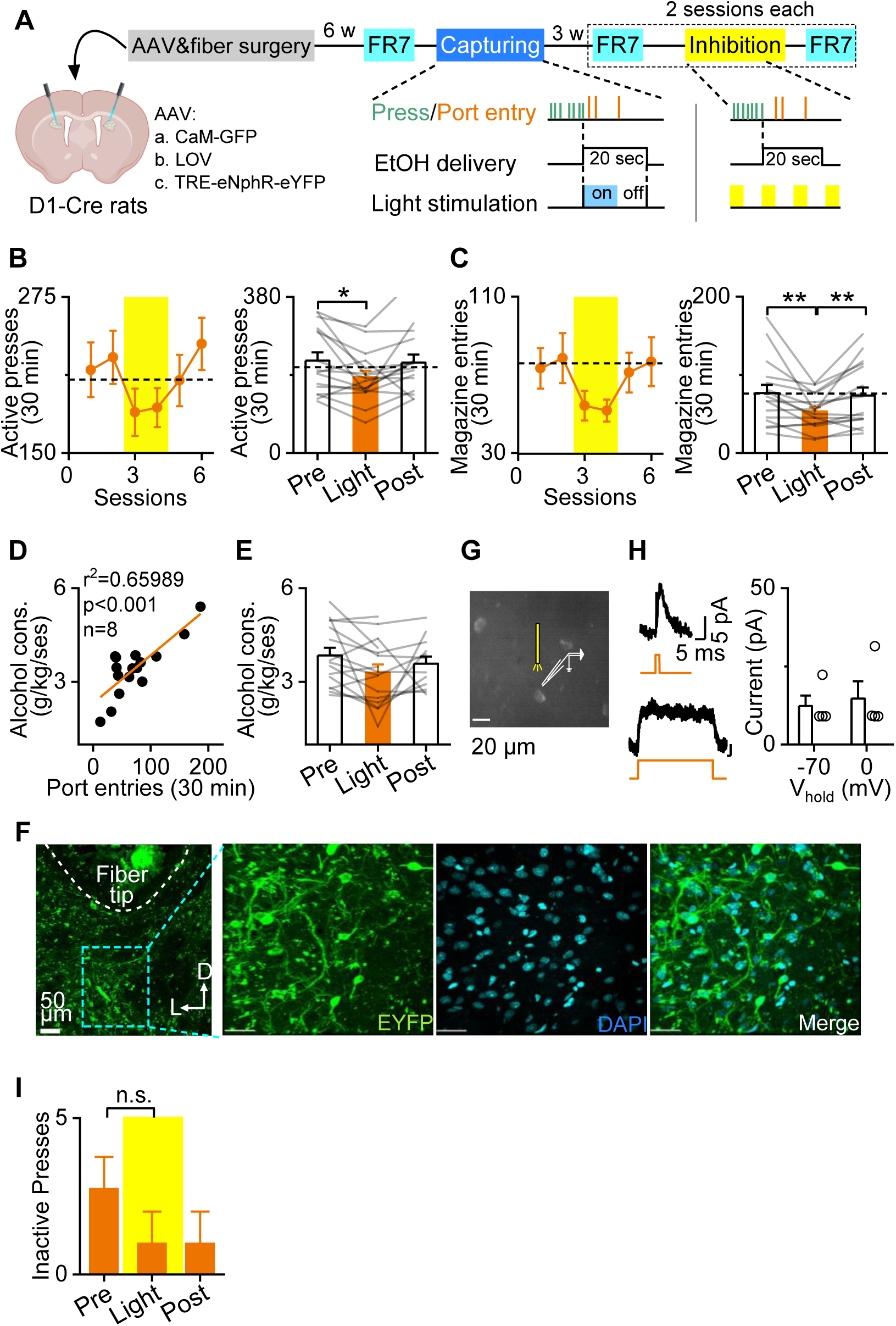
Optogenetic inhibition of magazine-entry-tagged neurons reduces subsequent magazine entries and lever presses. ***A,*** Experimental timeline; rats were infused with FLiCRE AAVs driving the expression of eNphR-eYFP. The animals were trained for alcohol operant self-administration (OSA) using a fixed ratio 7 (FR7) schedule. Blue light (470 nm) was delivered to tag magazine-entry-associated neurons during the first 10 seconds of a magazine entry. Three weeks later, yellow light (590 nm; every 2 seconds) was delivered during OSA to inhibit the magazine-entry-associated neurons. ***B,*** Time course (left) and bar graph (right) of active presses, depicting that optogenetic inhibition of magazine-entry-tagged neurons suppresses active presses. *p < 0.05, one-way RM ANOVA. n = 7 rats. ***C***, Time course (left) and bar graph (right) of magazine-entries, depicting that optogenetic inhibition of magazine-entry-tagged neurons suppresses magazine entries. *p < 0.05, **p < 0.01, one-way RM ANOVA. n = 4 rats. ***D,*** Correlation of the number of magazine entries and alcohol consumption in g/kg/ses. ***E***, Optogenetic inhibition of magazine-entry-tagged neurons suppressed alcohol consumption. One-way RM ANOVA. n = 4 rats. ***F***, Confocal imaging to confirm eNpHR-EYFP expression in the DMS underneath the tip of the optical fiber. Sections were stained with DAPI. D, dorsal; L, lateral. ***G***, Sample imaging of whole-cell recording in eNphR-positive neurons from behaviorally tested animals. ***H***, Yellow light stimulation of eNphR positive neurons induced direct hyperpolarization. ***I,*** There were no statistically significant changes in inactive lever presses before, during, or after light stimulation. n.s.=not significant, one-way RM ANOVA. n=4 rats.

Before beginning manipulations, we waited for three weeks to permit the expression of eNphR in the tagged neurons. Next, after two stable baseline sessions of OSA, we delivered yellow light (590 nm, two seconds on and two seconds off) to inhibit the magazine-entry-recruited/activated neurons during normal trials of OSA (Fig. 2A). We discovered that the inhibition of these neurons significantly decreased lever presses (Fig. 2B, (left) F_(4, 61)_ = 2.660, *p* < 0.05, (right) F_(2, 30)_ = 4.736, *p* < 0.05) and magazine entries (Fig. 2C, (left) F_(3, 47)_ = 3.109, *p* < 0.05, (right) F_(2, 25)_ = 8.183, *p* < 0.01). These reductions recovered following the removal of yellow light stimulation. In addition, we found that alcohol consumption, which was significantly correlated with magazine entries (Fig. 2D, *R^2^*=0.660, *p* < 0.001), decreased with the inhibition of magazine-entry-tagged neurons (Fig. 2E, *F*_(2, 22)_ = 3.285, *p* = 0.056). These findings indicate that alcohol-taking-associated neurons mediate both alcohol-seeking and -taking behaviors during OSA.

Six weeks following the optogenetic manipulations, we sacrificed animals and prepared coronal sections containing the DMS. We conducted confocal imaging to assess the expression of eNpHR. We found a large number of neurons in the DMS expressing eNpHR-EYFP underneath the tip of the optical fiber (Fig. 2F). Additionally, we prepared DMS slices and conducted whole-cell patch-clamp recording in eNpHR-positive neurons (Fig. 2G). We found that the delivery of yellow light-induced hyperpolarization of these neurons (Fig. 2H). These results validated the use of TRE-eNpHR-eYFP to inhibit the trapped neurons, permitting the FLiCRE manipulations. Finally, we ensured that there were no significant decreases in inactive lever presses (Fig. 2I, F_(2,6)_=1.138, *p*=0.381), showing that the changes in active lever presses weren’t due to motor deficits.

### Inhibition of magazine-entry-tagged neurons facilitates extinction learning

The neurons that contribute to OSA of alcohol may be targeted during the extinction of alcohol memory. Extinction training is hypothesized to suppress the original drug-related memory by inhibiting ensemble cells encoding drug-seeking and -taking behaviors. Given that alcohol-taking neurons mediate both alcohol seeking and taking during OSA, we next examined whether optogenetic inhibition of OSA-tagged magazine-entry neurons can further facilitate extinction learning. To do this, we delivered yellow light to inhibit these neurons during the first three extinction sessions of the same rats used in the prior experiment.

Inhibition of these OSA-tagged magazine entry neurons during extinction facilitated the extinction of alcohol-seeking (Fig. 3A, C-D; F_(1,7)_ = 8.5, p < 0.05), with the group undergoing inhibition having significantly fewer active presses than the control group (Fig. 3B, *t*_(5)_ = 2.913, p < 0.05). These results suggest that optogenetic inhibition of OSA-related alcohol-taking neurons enhances the effects of extinction training. This further indicates that extinction learning reduces the activity of these OSA-tagged magazine-entry neurons.

**Figure 3.**
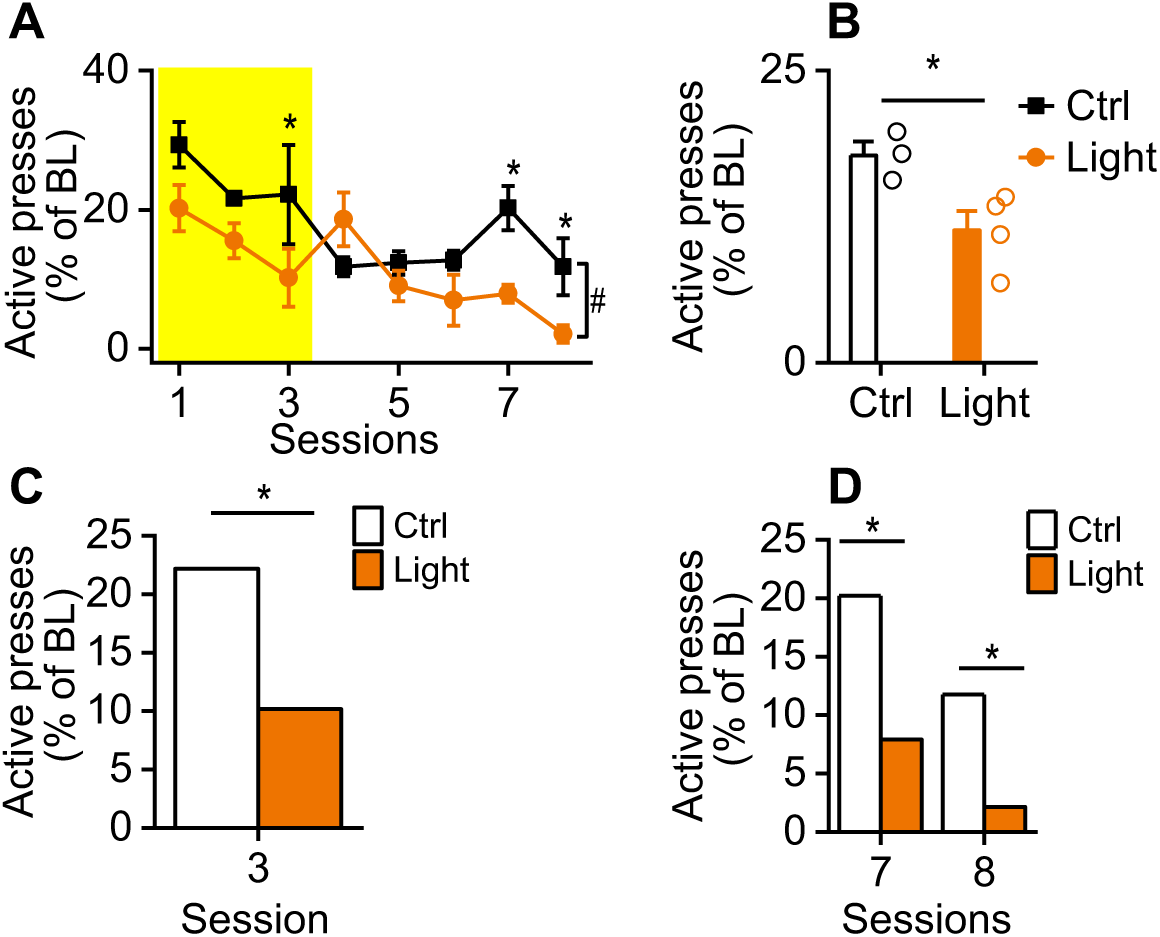
Optogenetic inhibition of magazine-entry-tagged neurons facilitates extinction. ***A,*** Time course of active presses following optogenetic inhibition of magazine-entry-tagged neurons (shown by yellow in the graph) during extinction, normalized to the baseline responding in alcohol OSA. *^#^p* < 0.05; \**p* < 0.05, Two-way RM ANOVA. n = 3 (ctrl) and 4 (light). Control (Ctrl) animals were infused with TRE-EYFP. ***B,*** Bar graph of active presses during versus after optogenetic inhibition of magazine-entry tagged neurons during extinction compared to the control group. \**p* < 0.05, unpaired T-test. n = 3 (ctrl) and 4 (light). ***C,*** Bar graph of active presses during optogenetic inhibition of magazine-entry tagged neurons during extinction compared to the control group. \**p* < 0.05, Two-way RM ANOVA. n = 3 (ctrl) and 4 (light). ***D,*** Bar graph of active presses after optogenetic inhibition of magazine-entry tagged neurons during extinction compared to the control group. \**p* < 0.05, Two-way RM ANOVA. n = 3 (ctrl) and 4 (light).

### Inhibition of magazine-entry-tagged neurons during reinstatement decreases alcohol-seeking behaviors

Reinstatement is known to reactivate neurons recruited by alcohol-seeking and -taking behaviors during OSA training (Bobadilla et al., 2020). Because of this, we examined if optogenetic inhibition of OSA-tagged magazine-entry neurons affected alcohol-seeking and - taking behaviors during reinstatement. We found that inhibition of alcohol-taking neurons significantly decreased the reinstatement of alcohol-seeking behaviors (Fig. 4A, *t*_(11)_ = 3.771, P < 0.01). The group that underwent inhibition maintained similar levels of lever-pressing behaviors as seen during extinction, whereas the control group had significantly increased lever-pressing behaviors compared to extinction levels of responding (Fig. 4B (left), *t*_(12)_ = −3.406, P < 0.01, (right), *t*_(10)_ = 0.834, P > 0.05; Fig. 4C, *t*_(11)_ = 2.842, P < 0.05). These results demonstrate that the inhibition of OSA-tagged alcohol-taking neurons can suppress the reinstatement of alcohol-seeking behaviors.

**Figure 4.**
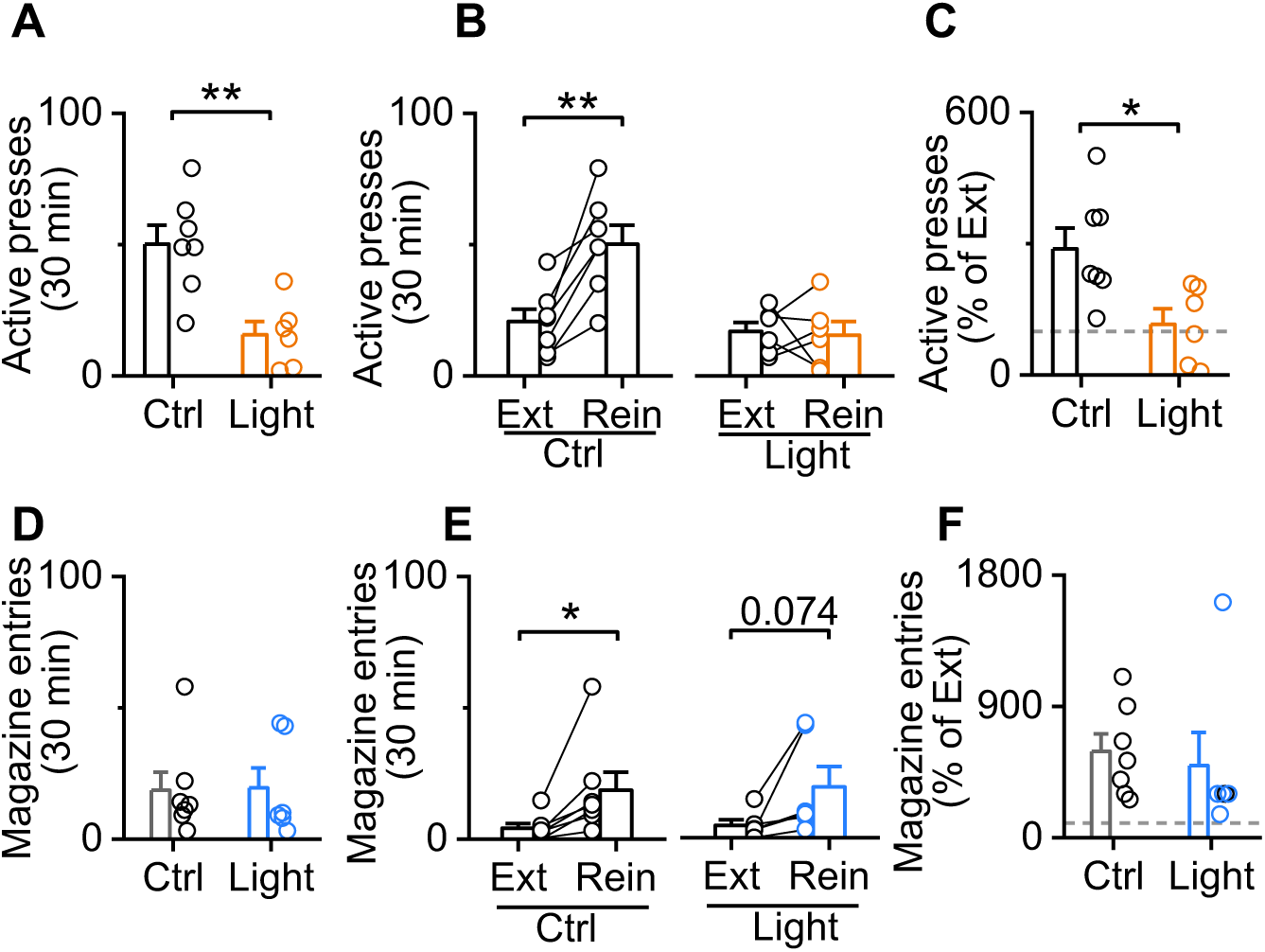
Optogenetic inhibition of magazine-entry-tagged neurons suppressed the reinstatement of alcohol seeking. ***A***, Optogenetic inhibition of magazine-entry-tagged neurons suppressed active lever presses during reinstatement. \*\**p* < 0.01, unpaired T-test. n = 7 (Ctrl) and 6 (Light). ***B***, Optogenetic inhibition of magazine-entry-tagged neurons suppressed active lever presses to levels similar to extinction responding whereas control animals had significantly increased lever presses. \*\**p* < 0.01, unpaired T-test. n = 7 (Ctrl) and 6 (Light). ***C***, Optogenetic inhibition of magazine-entry-tagged neurons suppressed active lever presses during reinstatement when normalized to extinction levels of responding. \**p* < 0.05, unpaired T-test. n = 7 (Ctrl) and 6 (Light). ***D***, Optogenetic inhibition of magazine-entry-tagged neurons did not affect magazine-entries during reinstatement. Unpaired T-test. n = 7 (Ctrl) and 6 (Light). ***E***, Optogenetic inhibition of magazine-entry-tagged neurons did not suppress magazine entries to levels similar to extinction responding. \**p* < 0.05, unpaired T-test. n = 7 (Ctrl) and 6 (Light). ***F*,** Optogenetic inhibition of magazine-entry-tagged neurons did not suppress magazine entries during reinstatement when normalized to extinction levels of responding. Unpaired T-test. n = 7 (Ctrl) and 6 (Light).

Interestingly, however, inhibition of OSA-tagged magazine entry neurons did not significantly decrease levels of magazine entries in reinstatement (Fig. 4D, *t*_(11)_ = 0.930, P > 0.05). Additionally, there was an increase in magazine-entry behaviors compared to extinction levels of responding in both the group that underwent inhibition and the control group (Fig. 4E, (left) *t*_(12)_ = −2.013, P < 0.05, (right) *t*_(10)_ = −1.856, P > 0.05; Fig. 4D, *t*_(11)_ = 0.394, P > 0.05). These results suggest that despite significantly suppressing the reinstatement of alcohol-seeking behaviors, inhibition of OSA-tagged alcohol-taking neurons do not have as strong an effect in the reinstatement of alcohol-taking behaviors.

## DISCUSSION

In this study, we utilized FLiCRE during alcohol OSA to precisely capture and manipulate striatal ensembles activated during alcohol-taking behaviors. We demonstrated that the optogenetic inhibition of alcohol-taking (magazine entry) neurons suppressed both alcohol-seeking and -taking behaviors during OSA. We also determined that the suppression of OSA-tagged alcohol-taking neurons facilitated the extinction of alcohol-seeking behaviors. Finally, we demonstrated that suppressing OSA-tagged alcohol-taking neurons suppressed the reinstatement of alcohol-seeking. These results suggest that alcohol-taking neurons within the DMS encode alcohol-seeking memories to maintain operant alcohol self-administration and drive relapse.

The majority of prior studies looking at the role of the DMS in AUD have focused on alcohol-seeking behaviors (Corbit et al., 2012, Barker et al., 2015). However, the studies that have looked at alcohol-taking behaviors showed that the DMS also plays a critical role in consumption (Chen et al., 2011, Wang et al., 2010). Our findings are in line with previous studies which indicated that alcohol consumption alters glutamatergic transmission in the DMS, leading to future alcohol-seeking behavior (Roltsch Hellard et al., 2019, Ma et al., 2017, Wang et al., 2012). These findings provide further evidence supporting the idea that the DMS plays a role in alcohol consumption behaviors and indicate that the neurons implicated in alcohol-taking behaviors also contribute to alcohol-seeking behaviors.

We showed that inhibition of OSA-tagged alcohol-taking neurons facilitated the extinction of alcohol-seeking behaviors, suggesting that these neurons are implicated in the regulation of alcohol seeking during extinction. It is highly likely that the ensembles encoding alcohol seeking and taking behaviors have strong overlap, which could cause manipulation of one to impact the other. Since both of the ensembles associated with seeking and taking behaviors mediate alcohol memory, it is likely that the inhibition of one will facilitate the extinction phenotype. Additionally, in the extinction operant set-up, there is no alcohol received following alcohol-seeking behaviors. Thus, it is possible that the same ensemble of neurons encodes the entire action sequence seen in extinction. Likely due to the nature of extinction paradigms, wherein no alcohol is delivered, most studies looking at the extinction of alcohol OSA have focused on alcohol-seeking rather than -taking behaviors. However, one study found that administration of Naltrexone, an opioid receptor antagonist which decreases alcohol consumption (Rösner et al., 2010), also decreased alcohol seeking during extinction in baboons (Kaminski et al., 2012). A similar study examined the effects of Baclofen, a GABA-B receptor agonist that reduces craving and alcohol consumption in patients (Rozatkar et al., 2016), and found that it also decreased seeking responses in extinction. These findings lend credence to the idea that alcohol consumption behaviors play a critical role in the extinction of alcohol seeking. Subsequently, future research should aim to examine these behaviors more in depth to better determine the specific role of consumption behaviors in the extinction of alcohol seeking.

Interestingly, although reinstatement is thought to reactivate neurons recruited by both alcohol-seeking and -taking behaviors during OSA training, inhibition of OSA-tagged alcohol-taking neurons suppressed the reinstatement of alcohol-seeking behaviors but did not significantly suppress the reinstatement of alcohol-taking behaviors. These results suggest that the ensembles underlying alcohol-taking behaviors play a critical role in the extinction and reinstatement of alcohol-seeking behaviors, but not necessarily for the reinstatement of alcohol-taking behaviors.

A primary strength of FLiCRE is that it is an activity-dependent labeling system which permits high temporal resolution due to its reliance on calcium signals (Kim et al., 2020). Other trapping mechanisms, such as those relying on immediate early gene (IEG) expression, lack this advantage, taking hours to mark neuronal activity (Casey et al., 2013). FLiCRE, on the other hand, provides a minute-timescale sensitivity. In addition, FLiCRE offers a higher calcium sensitivity than FLARE, a predecessor which also offered a light- and calcium-based tagging approach for higher temporal resolution (Wang et al., 2017). FLiCRE requires only 30 seconds of elevated calcium and light for substantial tagging, whereas FLARE was shown to require closer to 15 minutes (Kim, 2023). Additionally, FLiCRE had a tagging resolution of 1 minute, suggesting that neurons would only be tagged if they were both active and had light delivered within one minute. These benefits permit the analysis of short-term behaviors and the differentiation of the neuronal ensembles mediating them. Alcohol-seeking and -taking are two such behaviors, particularly in rodent models of OSA, with there being only a brief period between lever presses and alcohol delivery. Subsequently, FLiCRE was an excellent tool to examine the differences in cellular substrates underlying seeking and taking behaviors. The successful use of FLiCRE in our study demonstrates that this methodology offers a promising future for the study of neuronal ensembles involved in different time-sensitive behaviors.

Despite the benefits of FLiCRE, there are several limitations to the method that must be considered. First, this system requires three viruses to work together. Subsequently, it is challenging to ensure that the expression level is at the correct ratio for these three proteins; reaching the correct ratio is necessary to have the correct levels of the protease LOV expressed. Although there have been efforts to reduce the number of necessary viruses with the similar trapping mechanism single-chain FLARE (scFLARE) (Sanchez et al., 2020), it may still be difficult to completely resolve this issue. Future studies may want to assess different methods to improve this area. Additionally, because the light stimulation required in FLiCRE is restricted to a specific area, this method is limited to a specific brain region. This method differs from other trapping mechanisms such as Fos/ArcTRAP, which can label the whole brain. To solve this issue, an interesting prospect may be to employ the use of focused-ultrasound with liposomal-based particles containing chemiluminescent compounds for non-invasive optogenetic stimulation (Wang et al., 2023). With this method, ultrasound stimulation promotes these particles to produce blue light, offering a potential method to optogenetically target larger brain regions due to not requiring the implantation of an optical fiber.

Several behavioral treatment methods for AUD rely on the use of extinction-based mechanisms, aiming to prevent relapse (as can be measured using reinstatement). Additionally, as both alcohol-seeking and -taking behaviors contribute to the pathophysiology of AUD, a greater understanding of the neural ensembles underlying these behaviors is critical to the development of future therapeutic strategies. Our use of FLiCRE to differentiate between the ensembles mediating different AUD behaviors is novel and promising for future research aiming to dissect these behavioral circuits. Overall, our study found that inhibiting magazine-entry tagged OSA neurons in the DMS led to reduced alcohol-taking and -seeking behaviors, facilitated the extinction of seeking behaviors, and prevented the reinstatement of seeking behaviors. This comprehensive approach reveals the involvement of specific neuronal ensembles in alcohol-seeking and -taking behaviors, shedding light on potential mechanisms underlying AUD and the use of potential methods for further study.

## ACKNOWLEDGEMENTS

This work was supported by NIH grants: U01AA025932 (to JW), R01AA021505 (to JW), R01AA027768 (to JW), R01AA030293 (to JW); by an X-Grant from the Presidential Excellence Fund at Texas A&M University (to JW); and by McGovern Fellowship from Texas Research Society on Alcoholism (to XX).

## AUTHOR CONTRIBUTIONS

JW conceived, designed, and supervised all the experiments in the study. VV, XX, JW, and ZH wrote and edited the manuscript. ZH and XX designed and performed electrophysiological experiments and analyzed the data. XX designed and conducted behavioral experiments. XX and VV analyzed the data from the behavioral experiments. XX conducted histology experiments and analyzed the data. XX, HL, EB, RS, and YZ developed the neuronal culture and AAV packing protocols. DW bred transgenic animals and conducted genotyping.

## DECLARATION OF INTERESTS

The authors declare no competing interests.

## Notes

### Competing Interest Statement

The authors have declared no competing interest.

